# Pair Matcher (*PaM*): fast model-based optimisation of treatment\case-control matches using demographic and genetic data

**DOI:** 10.1101/145177

**Authors:** Desmond M Ryan, Eran Elhaik

**Affiliations:** University of Sheffield, Department of Animal and Plant Sciences, Sheffield, UK S102TN; University of Sheffield, INSIGNEO Institute for In Silico Medicine, Sheffield, UK S13JD

**Keywords:** population structure, population stratification, clinical trials, randomised controls, Principal Component Analysis (PCA), association studies

## Abstract

In clinical trials, individuals are matched for demographic criteria, paired, and then randomly assigned to treatment and control groups to determine a drug’s efficacy. The successful completion of pilot trials is a prerequisite to larger and more expensive Phase III trials. One of the chief causes for the irreproducibility of results across pilot to Phase III trials is population stratification bias caused by the uneven distribution of ancestries in the treatment and control groups. Pair Matcher (*PaM*) addresses stratification bias by optimising pairing assignments *a priori*- and\or *posteriori* to the trial using both genetic and demographic criteria. Using simulated and real datasets, we show that *PaM* identifies ideal and near-ideal pairs that are more genetically homogeneous than those identified based on racial criteria or Principal Component Analysis (PCA) alone. Homogenising the treatment (or case) and control groups can be expected to improve the accuracy and reproducibility of the study. *PaM*’s ability to infer the ancestry of the participants further allows identifying subgroup of responders and developing a precision medicine approach to treatment. *PaM* is simple to execute, fast, and can be used for clinical trials and association studies. *PaM* is freely available via R scripts and a web interface.

## Introduction

It is well recognized that pharmaceutical research and development (R&D) is in a crisis. The number of new drugs approved per billion US dollars spent on R&D has halved roughly every 9 years since 1950, falling around 80‑fold in inflation-adjusted terms (Scannell et al. 2012) as spending in the industry has inflated to an average of ~$5.8 billion per drug in 2011 compared to $1.3 billion per drug in 2005 (Roy 2012). Clinical trials are typically divided into several phases, the earliest of which evaluate the safety, dosing, and side effects of the drug (Pilot or Phase I trials). The latter phases test the drug’s efficacy compared to a placebo or other treatments in a randomised trial setting and require assessing tens, hundreds (Phase II trials), and eventually tens of thousands (Phase III trials) of volunteers over a long time period to prove that there is substantial evidence for the clinical benefit of the drug. Under the Federal Food, Drug, and Cosmetic (FD&C) Act, the FDA has considerable discretion to determine what constitutes “substantial evidence.” The FDA typically requires two large multiyear Phase III trials showing with 95% statistical certainty that a drug meets its tested aims of clinical benefit. Only one in 12 drugs that enter human clinical trials end up gaining approval from the FDA (Roy 2012). While the drivers behind this crisis are still under debate (Scannell et al. 2012), it is acknowledged that one of the biggest drivers to the increase in R&D costs is the regulatory process governing Phase-III clinical trials of new pharmaceuticals (Roy 2012). Since it is unlikely to expect a relaxation of the regulatory environment (Scannell et al. 2012), it is important to understand why randomised control trials may be more successful in smaller than larger trials.

Matching treatment (or case) with control groups is the most elementary and critical part of any trial (or study). Mismatched groups introduce genetic heterogeneity that may obscure performance of the trialed drug due to genetic predisposition to response, and result in reduced reproducibility between different cohorts (Scannell et al. 2012). Currently, individuals are matched based on demographic criteria (e.g., age, gender, self-reported “race”) and then randomly assigned to treatment and controls groups. It is well acknowledged that due to the significant heterogeneity among humans, demographic-based matching alone is inadequate (e.g., Toobert et al. 2010). Trials are thus vulnerable to ‘stratification bias,’ a result of differences in genetic ancestry between individuals, which is not factored in when trial participants are grouped based on demographics alone. This undetected bias may contribute to biased interpretation of trial results due to lack of genetic information that may confound interpretation, leading to alterations in numbers of false negative or false positive results, with subsequent financial and patient health consequences (Figure 1a). In large groups the stratification bias may be less pronounced, however it is practically unavoidable in the case of rare diseases due to the difficulties in recruiting genetically homogeneous participants (Yusuf and Wittes 2016). Crucially, this bias is more severe in small cohorts leading to an applied misinterpretation of the drug’s efficacy that will be difficult to replicate in larger trials.

**Figure 1a.**
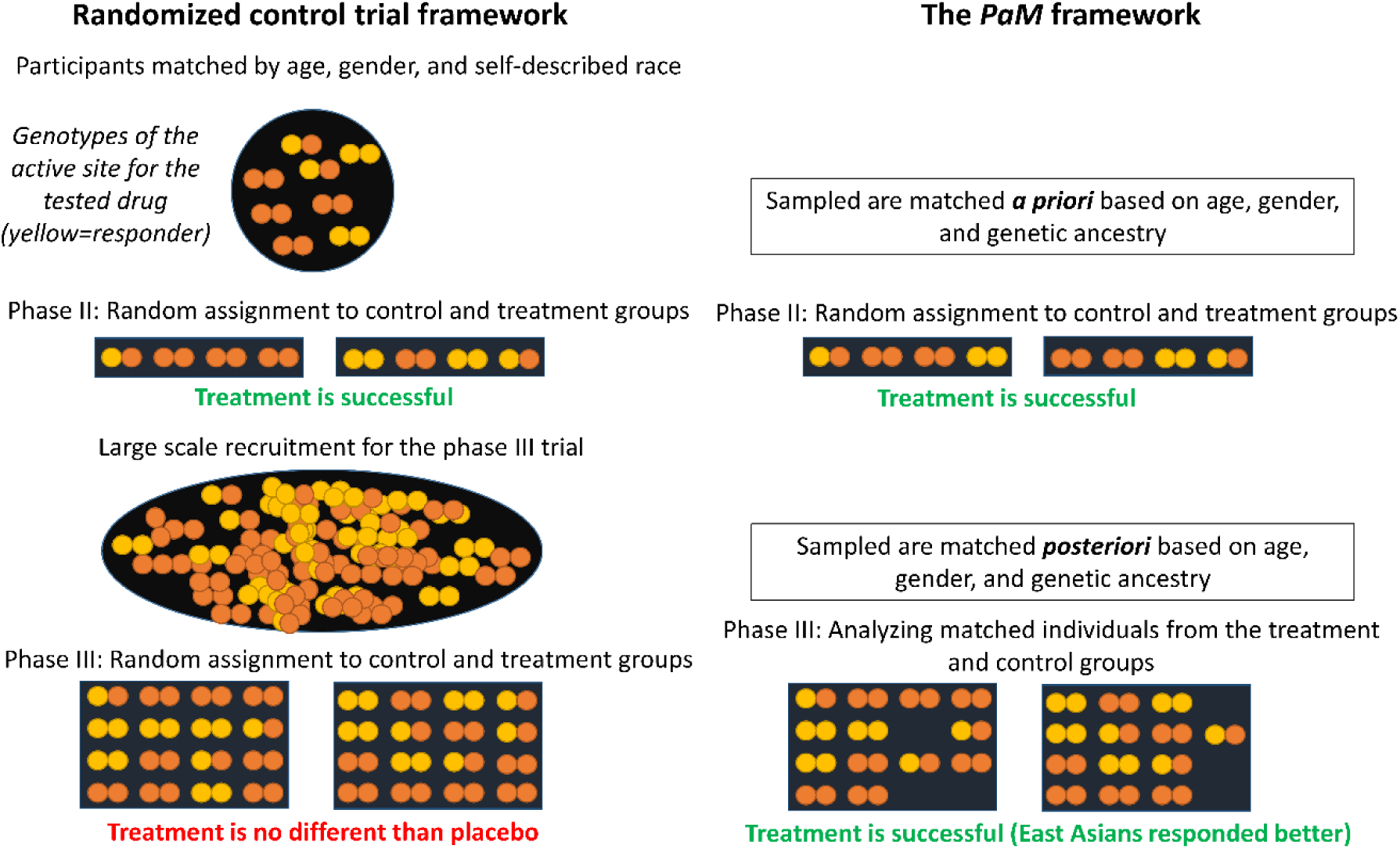
Comparing the framework of a standard controlled trial and with *PaM*. First, a cohort of age, gender, and race-matched participants is identified and randomly divided into control and treatment groups. Unknown to the tester, participants with yellow alleles respond better to the drug. The uneven distribution of this allele in the Phase III trial deemed the treatment ineffective. *PaM* homogenizes the samples both *a priori* and *posteriori* to the trial. After analysing *PaM*’s generated cohort the treatment is considered successful. Moreover, by analyzing *PaM*’s records, it appears that East Asians are better responders, allowing further development of a personalized medicine approach to the treatment.

Population stratification can be addressed by optimising the treatment-control matches *a priori* or\and *posteriori* to the trial using a variety of tools applied to the genotype data and selecting matched pairs for downstream analyses. Due to the historically high cost of genotyping and sequencing, *a priori* methods rely heavily on demographic-based matching criteria followed by statistical corrections made *posteriori*, if at all. *A priori* methods have long been considered biased, inaccurate, and unhelpful (De Bono 1996; McAuley et al. 1996; Fustinoni and Biller 2000) due to their reliance on self-reported “race” (“Africans,” “Asians” and “European-Americans” or “Whites”) or regional similarity, which does not eliminate the bias (Campbell et al. 2005; Wang, Localio, and Rebbeck 2006; Chikhi et al. 2010; Elhaik et al. 2014; Yusuf and Wittes 2016). Unable to completely account for choices made at the *a priori stage, posteriori* methods may make simplifying, unrealistic, or problematic assumptions (Kimmel et al. 2007), particularly concerning population structure. For example, “genomic control” is a non-parametric method for controlling the inflation of test statistics due to population structure, which assumes that the inflation factor *λ* is constant throughout the tested region and that every SNP is in null-association with the trait (Devlin, Roeder, and Wasserman 2001). The genomic control method attempts to estimate the amount by which association scores are inflated by *λ.* In large-scale studies that evaluate heterogeneous regions this assumption does not hold and can lead to loss of power and control over the test level (Epstein, Allen, and Satten 2007; Bouaziz, Ambroise, and Guedj 2011). Even if an appropriate null distribution is obtained, this approach will not maximize power to detect true associations (Price et al. 2010). Other methods compute the principal components (PCs) of the genotype matrix and adjust the genotype vectors by their projections on the PCs (e.g., Price et al. 2006). However, linear projections cannot be assumed to sufficiently correct for the effect of stratification due to other unaccounted confounders (Kimmel et al. 2007). PCs ignore the complexity of population structure, are influenced by uneven sampling, and so have diminished sensitivity for handling individuals that are of mixed origins (McVean 2009; Yang et al. 2012; Elhaik et al. 2014; Lacour et al. 2015). Even newer tools (Epstein, Allen, and Satten 2007; Kimmel et al. 2007; e.g., Lacour et al. 2015) can make only basic assumptions concerning population structure, may ignore admixture, or do not fit a clinical trial.

Prompted by the high cost of clinical trials and the plummeting cost of genotyping (~$60/sample) and sequencing (projected to reach $100/sample), we propose accounting for population stratification *a priori* to the trial; matching samples by demographic and genetic criteria. We have developed the Pair Matcher (*PaM*) – A genetic-based tool that optimises pairing assignments *a priori*- and\or *posteriori* to the trial, and so allows trial designers to make informed decisions in real time (Figure 1a). *PaM* models individual genomes as consisting of gene pools (or admixture components) that correspond to their recent demographic history (Elhaik et al. 2014). *PaM* then matches individuals based on their age, gender, and the similarity of their admixture components using *PaM*_*simple*_ or *PaM*_*full*_. We first compared the accuracy of these algorithms and then the accuracy of the best performing algorithm to pairings made either at random, based on racial-criteria, or through PCA (e.g., Luca et al. 2008). Optimising the trial design can be expected to homogenize the treatment (or case) and control groups and improve the accuracy and reproducibility of the study. This can be expected to lower drug developmental costs and benefit the patients. An added advantage to *PaM* is the ability to infer the ancestry of the responders and develop a precision medicine approach to treatment.

## Results

To test the ability of *PaM* and various approaches to identify genetically homogeneous pairs, twenty-four simulated datasets with 480-500 predetermined pairs were created. Each individual was represented by nine admixture components. In the first eight datasets, consisting of 500 pairs, the genetic distances between the pairs increased by a fixed factor in each dataset starting at 0 (identical pairs) in Dataset 1 making it increasingly difficult to match the original pairs. Genetic heterogeneity was achieved by perturbing the admixture components by a fixed amount per dataset and normalizing the components to sum to 1 (Figure 2). The remaining datasets were created by randomly removing one (Datasets 9-16 *remove-1*) or twenty individuals (Datasets 17-24 *remove-20*) from Datasets 1-8. Each algorithmically matched pair could have obtained a top score of 10 if the two individuals had similar age and gender (one point) as well as similar admixture components (nine points).

Pairing approaches were compared over the number of *matched pairs* and the remaining *unpaired individuals* consisted of individuals without a pair as well as poorly scored pairs if threshold was used (Figure 1b). For the matched pairs we calculated the total genetic distance (GD) between the admixture components of the matched pairs, their total score (1 for matching age and gender and 1 for each matching admixture component), and the number of *misassigned pairs*, i.e., pairs that are not with their original partners. An optimal pairing solution for Datasets 1-8 would consist of 500 pairs with a GD of 0 between all pairs and a total score of 5,000 (top score of 10 for 500 pairs). No individuals would be left unpaired or misassigned.

**Figure 1b.**
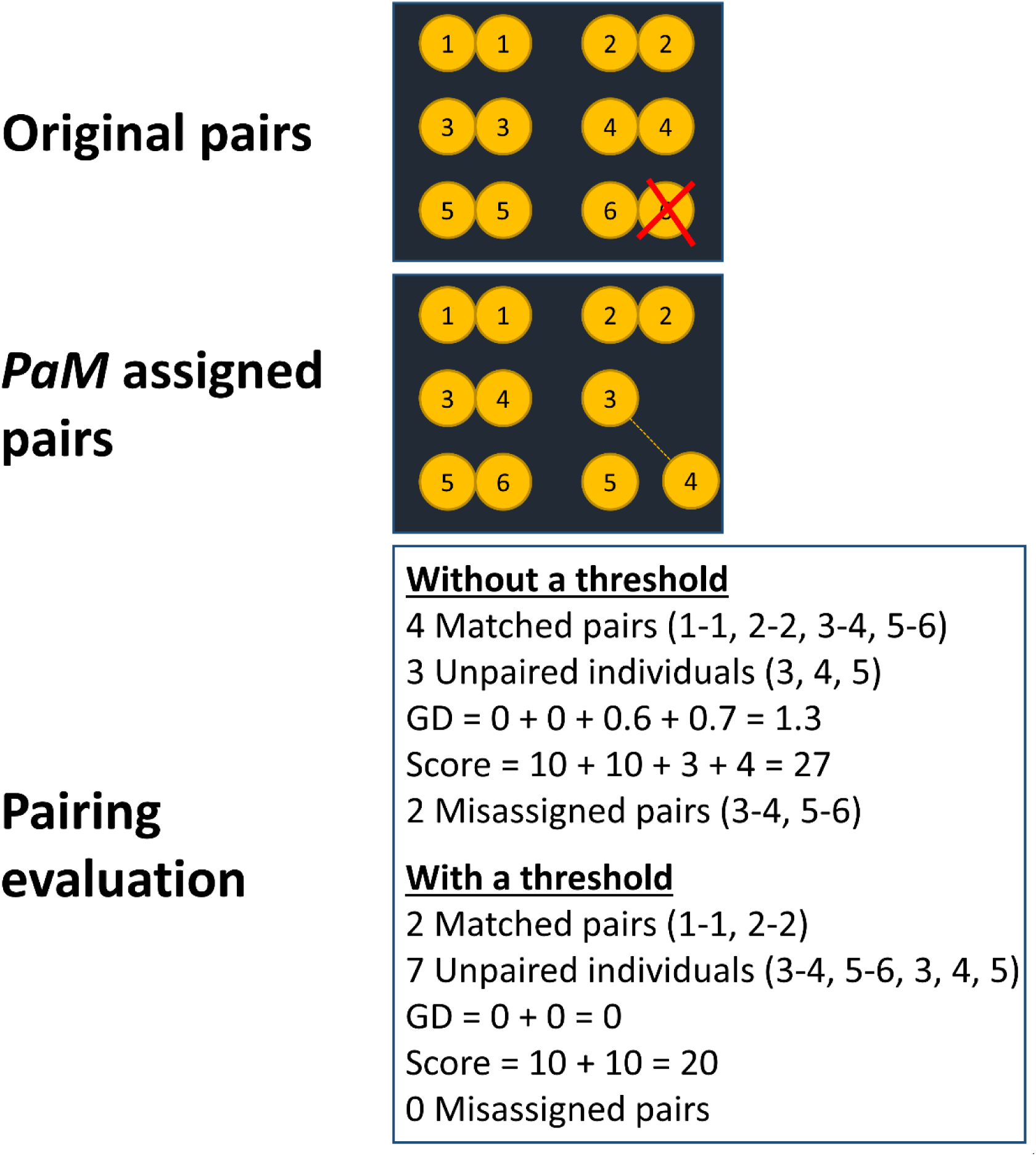
An illustration on how pairing solutions are evaluated. Consider an initial dataset with six genetically matched pairs. One individual was then randomly removed. An attempt was made to match the pairs using *PaM*_*simple*_ that connected 3-4 and *PaM*_*full*_. *PaM*’s pairing solutions with and without threshold are evaluated based on five criteria. The results differ based on whether or not a threshold was applied.

**Figure 2.**
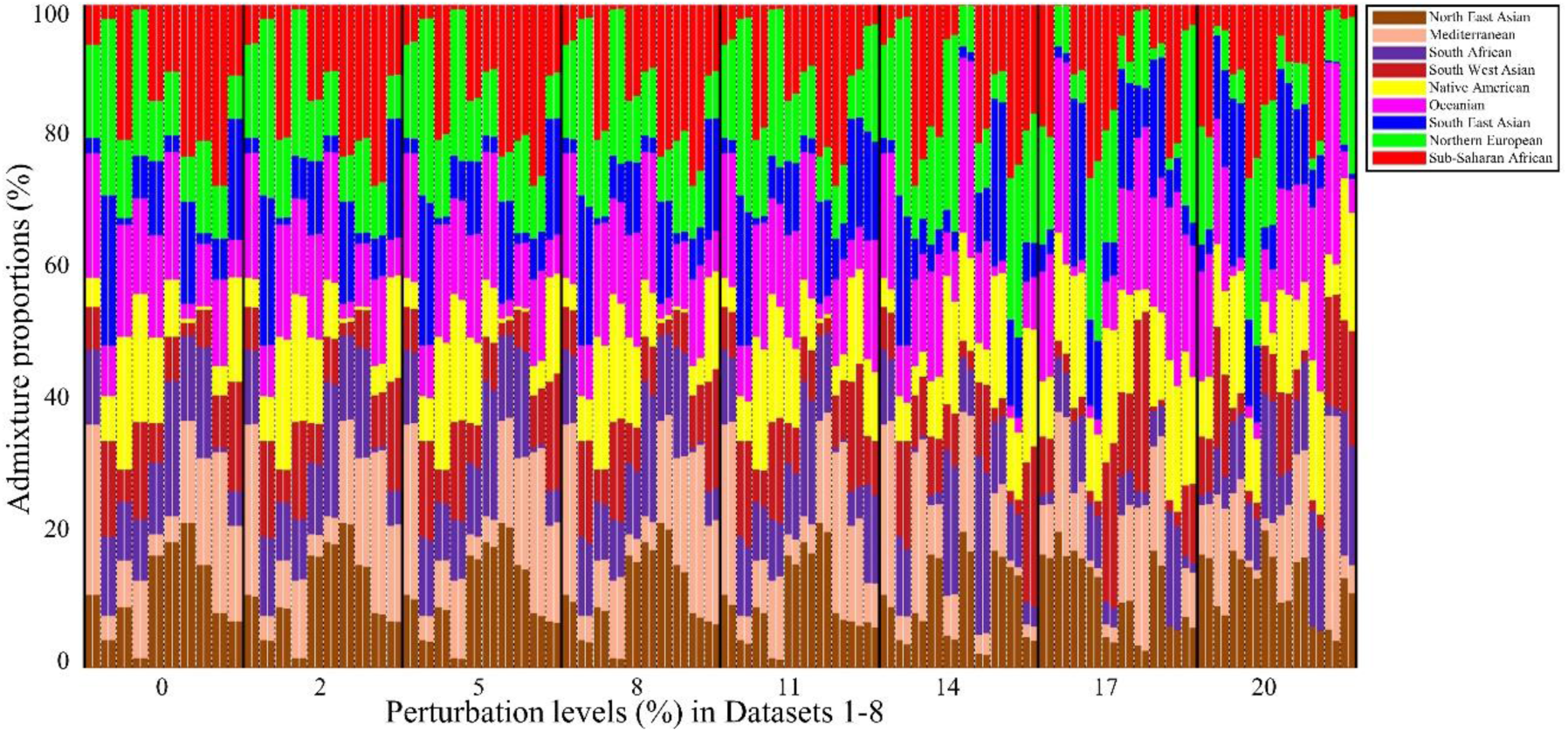
The admixture components of ten random pairs, shown consecutively, from Datasets 1-8. Each individual is represented by a vertical stacked column of colour-coded admixture components that reflects genetic contributions from the putative ancestral populations. The *x*-axis represents individual pairs from the unperturbed (0%) and perturbed (2-20%) datasets.

### Assessing PaM’s performances

We first evaluated the performances of *PaM*_*simple*_ without a threshold (no limit placed on the acceptable score of pairs) and with a threshold of 7 (necessitating the matching of age\gender and at least 6 admixture components) across all simulated datasets. As expected, when applying *PaM* without a threshold, the GD increased with increasing perturbation or heterogeneity while the score decreased, and the number of misassigned pairs increased (Figure 3, Supplementary Table 1). However, despite the perturbation and removal of individuals, most of the original pairs (80-100%) were correctly identified, particularly in Datasets 1-16. The increase in the number of misassigned pairs is related to how *PaM*_*simple*_ searches for an optimum solution. *PaM*_*simple*_ selects a case index *i* (row *i* of the GD matrix) and finds the best match for this index by locating the minimum GD in the row, corresponding to the best possible match. This, however, does not necessitate an ideal match considering all other individuals, some of whom are best left unpaired. Since *PaM*_*simple*_ does not leave any individual unpaired (for an even cohort), a poor pairing may create a “snow ball” effect triggering other poor pairings, resulting in an overall increased GD and reduced score for the final pairing solution.

**Figure 3.**
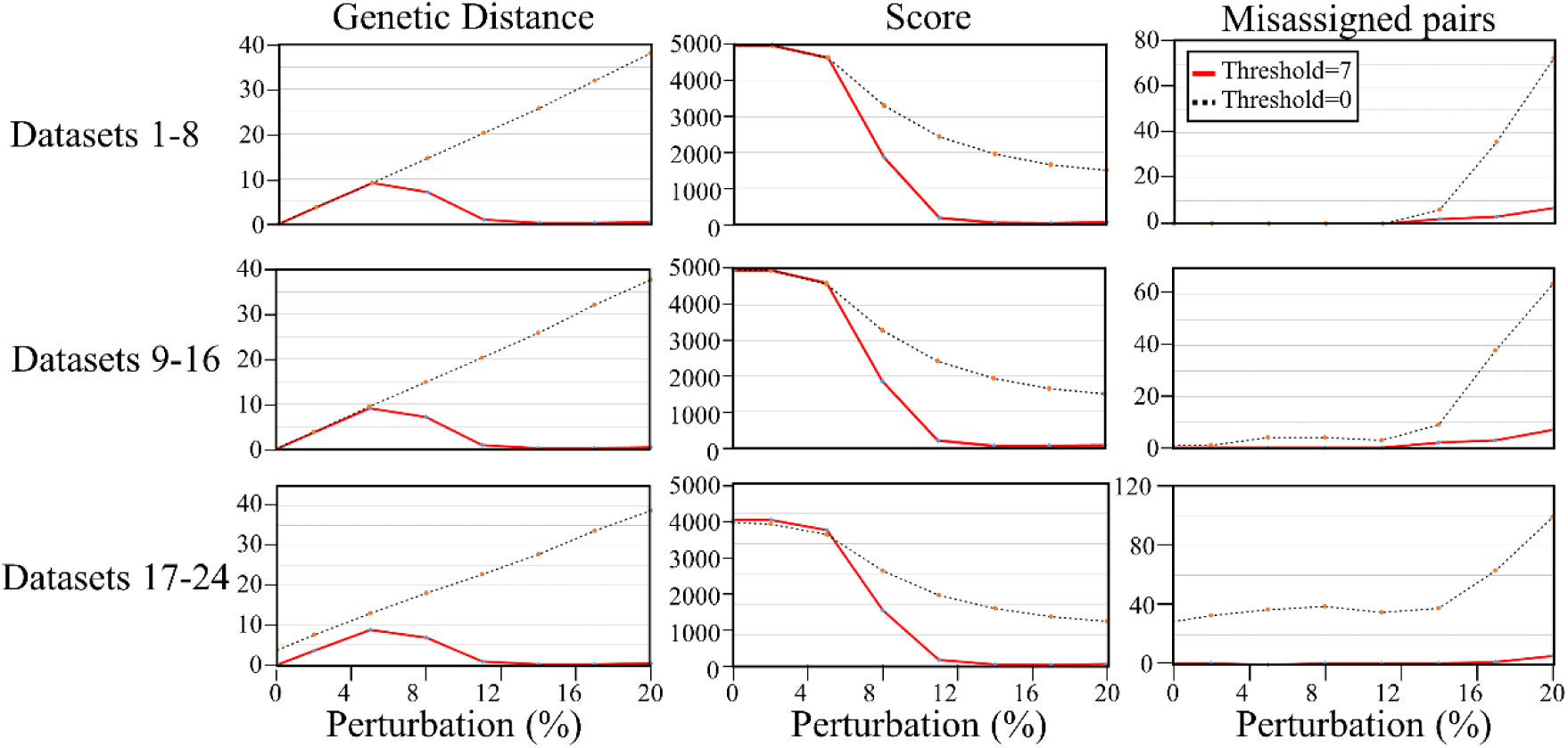
*PaM*_*simple*_ performances on 24 simulated datasets. Each row shows the results of eight perturbed datasets (full dataset, *remove-1*, and *remove-20*). From left to right, the columns show the total GD and pair score of the matched pairs, and the number of misassigned individuals. *PaM*_*simple*_ was applied without a threshold and with a threshold of 7.

We addressed this problem by applying a threshold of 7 on the match score. Under these settings, the GD curve decreased sharply at a perturbation level of ~5%; identifying genetically homogeneous pairs, despite the increased perturbation, and discarding genetically mismatched pairs. That the score decreased is due to the growing number of unpaired individuals, which represents the conservative choice on what is considered an acceptable pair. The trade-off to the low GD of acceptable pairs is that more individuals are left unpaired due to their low pairing score and are omitted from the total GD score (Figure 3, Supplementary Table 1). The advantage of applying a threshold is that it reduces the number of misassigned pairs by allowing only pairs with a high match score that are considered genetically homogeneous. This prevents the model from selecting pairs that satisfy the row GD minimum but do not have a favourable match score, thus avoiding the “snow ball” effect. Since the matrix is symmetric and each row has all possible pairing for each individual, individuals with a match score lower than the threshold are considered too genetically heterogeneous and placed on the unpaired list (Supplementary Table 2).

For Datasets 1-8, the best solution was obtained with a threshold at a perturbation level of 11%, where the GD was ~0 for all matched pairs, nearly half the individuals had an acceptable score, and the number of misassigned pairs was 0. We note that for heavier perturbations not all the misassignments are false positives since the perturbation created, by chance, more suitable pairing matches than the predefined ones. Considering the low GD between all pairs, the majority of matches were near-optimal ones even after removing individuals from the dataset. Interestingly, we observed a repeated single misassignment in most of the *remove-20* datasets (Supplementary Table 2). Examination of this unexpected misassignment showed it to be a pairing with a very low GD and a match score of 7 making it an acceptable assignment though not between the original partners, which could potentially be suboptimal.

There are two ways to address the vexing issue of ‘rogue’ misassignments. The first is to set a higher threshold and the second is to use *PaM*_*full*_, which carries out a more exhaustive pairing search by iteratively sorting (in ascending order) the cohort data by the admixture components. The pairing procedure for the cohort commences at a random row index (multiple times). This approach does not produce rogue misassignments and hence finds an optimum or near-optimum pairing solutions. The numerical results for the full, *remove-1*, and *remove-20* datasets using *PaM*_*full*_ are shown in Supplementary Tables S3 and S4. *PaM*_*full*_ results are similar to those of *PaM*_*simple*_, except that it does not allow the accidental misassignments (Supplementary Tables S4, perturbation <11%) observed with *PaM*_*simple*_ (Supplementary Tables S2). As before, the misassignments detected beyond the 11% threshold are due to the high similarity in admixture components in the post-perturbation stage and are not truly false positives. The cost of using *PaM*_*full*_ is an increased computation time, almost an order of magnitude greater than the *PaM*_*simple*_’*s* run time (~3 hours on a single core of an i5 processor). Due to its superior performances, the remaining of the analyses were done with *PaM*_*simple*_ with a threshold of 7.

### Comparing PaM’s performances to alternative methods

We next compared the performances of *PaM*_*simple*_ with two alternative methods, namely random assignment and three models of self-reported race (Figure 4). As expected, the GDs for the random assignment, where the age and gender matched but race was randomly determined, were much larger than the competing solutions. Correspondingly, the random assignment’s score is mostly lower than the alternative solutions. Nearly none of the pairs assigned at random were with their original counterparts.

**Figure 4.**
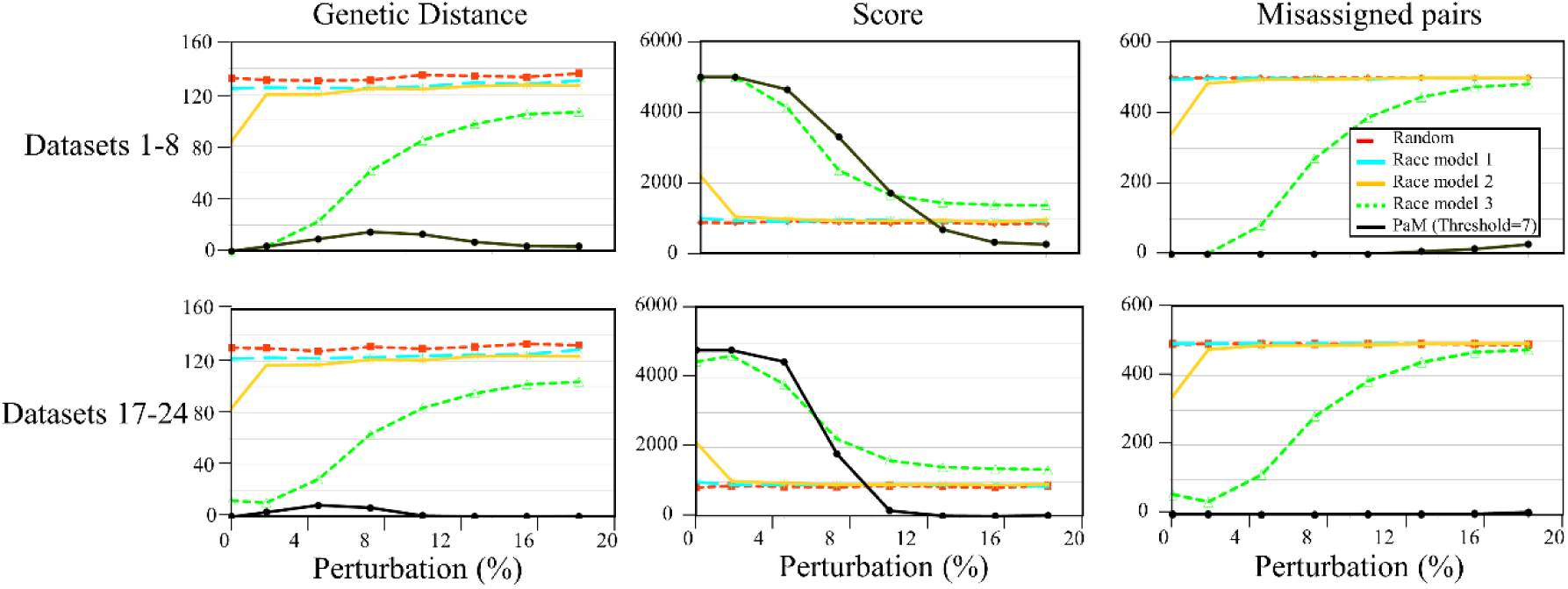
*PaM*_*simple*_ (threshold of 7) performances on 16 simulated datasets against two competing methods. Each row shows the results of the first eight datasets (row 1) and after omitting 20 individuals (*remove-20*, row 2). Columns show the total GD, pair score, and the number of misassigned individuals.

The first two self-reported race models (African, Asian, Latino, or White; African or non-African) perform only slightly better than the random assignment in terms of GD and the score. The results of the third model (mixtures of African, Asian, Latino, or White) are considerably better than the previous models or random assignments. This is to be expected, since this model can be considered a reduced form of *PaM*’s nine-admixture components model. Our results indicate that pairs obtained through standard demographics criteria (age, gender, and ethnicity) are as poor as those obtained at random.

### Comparing PaM’s performances with PCA

We next compared the performances of *PaM* with a PCA-based approach on the Genographic dataset consisting of worldwide individuals alongside 13 simulated Indian-British individuals. As before, *PaM*_*simple*_ was applied to the admixture components of all individuals. PCA was applied to the same dataset. Individuals were grouped into clusters based on the similarity of their first two eigenvectors using a *k*-means clustering approach. Individuals within each cluster were randomly paired.

We evaluated the homogeneity of the pairs inferred by *PaM* and PCA using both geographic and genetic distances, the latter calculated using identical by descent (IBS), as an impartial method. All of *PaM’s* inferred pairs had higher genetic similarity (i.e., smaller IBS distances) than PCA’s inferred pairs (Figure 5). We identified twelve IBS clusters (Figure 6) and divided all inferred pairs to “matches,” if individuals were in the same cluster, and “mismatches,” if otherwise. PCA pairs had 270 “matches” and 46 “mismatches” with mean distances of 1,042 km and 6,124 km, respectively and 10 unpaired individuals (Figure 7). *PaM* pairs had 284 “matches” and 17 “mismatches” with mean distances of 484 km and 557 km, respectively and 40 unpaired individuals. In comparison to PCA pairs *PaM* pairs had significantly smaller geographic distances between individuals regardless of the category (Kolmogorov-Smirnov goodness-of-fit test, *p-value* _(matches)_=2.74*10^−6^, *p-value* _(mismatches)_=3.58*10^−4^, *p-value* _(all)_=4.85*10^−10^). In one instance, *PaM* paired individual from Papua New Guinea with one from Peru, which yielded a geographic distance of over 13,000 km. However, Skogland et al. (2015) showed that some native American populations can trace their origins to Papua New Guinea, suggesting that *PaM’s* assignment was appropriate. This assignment was also made within the IBS’s cluster and classified as “match.” Overall *PaM* produced pairs that are genetically (Figure 5) and geographically (Figure 7) more similar than PCA.

**Figure 5.**
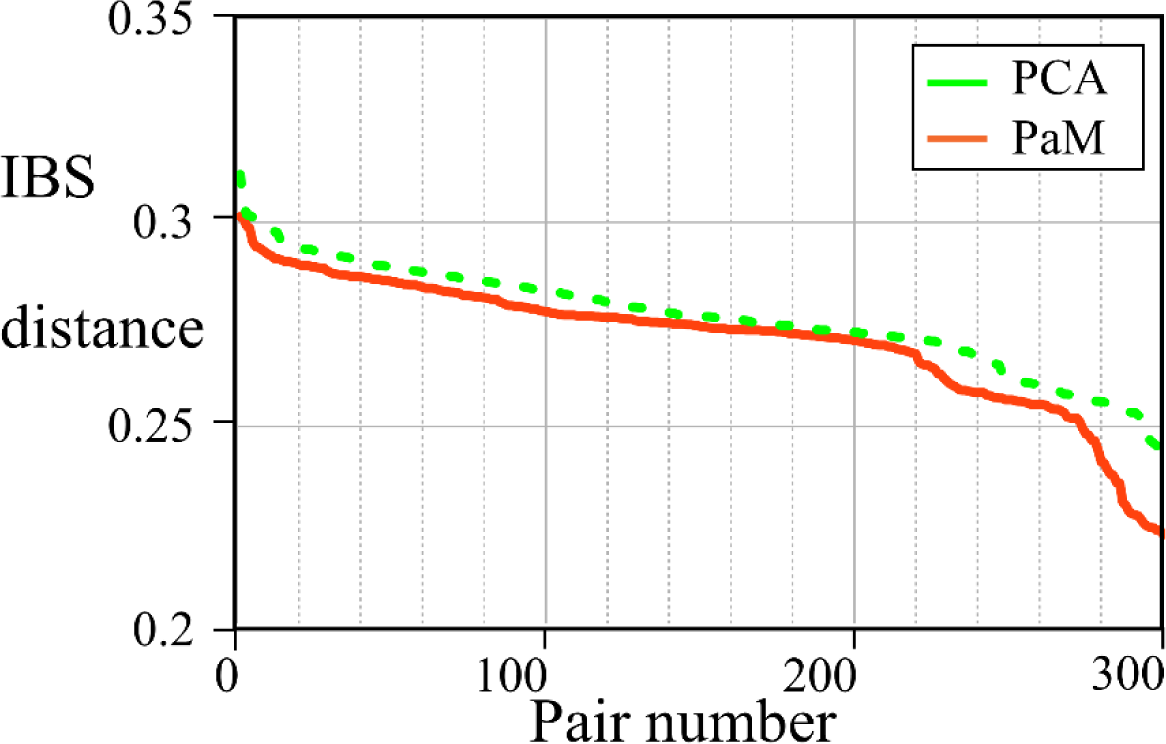
IBS distance between *PaM* and PCA inferred pairs.

**Figure 6.**
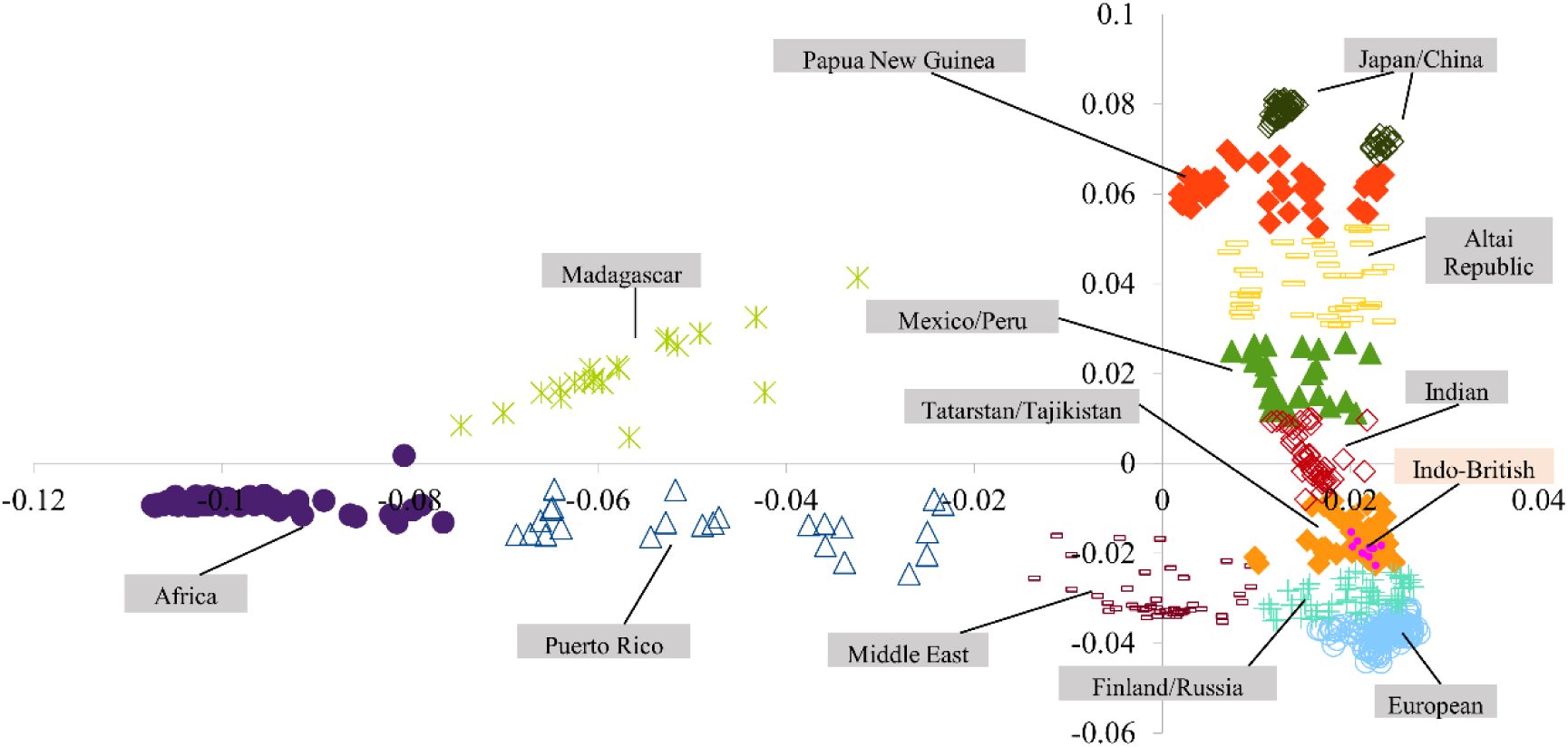
PCA plot of genetic diversity across the worldwide populations. The figure represents the genetic diversity seen across the populations considered, mapped onto a spectrum of genetic variation represented by two axes of variations corresponding to two eigenvectors of the PCA. Individuals from each population cluster are marked by region that are represented by unique shape and color.

**Figure 7.**
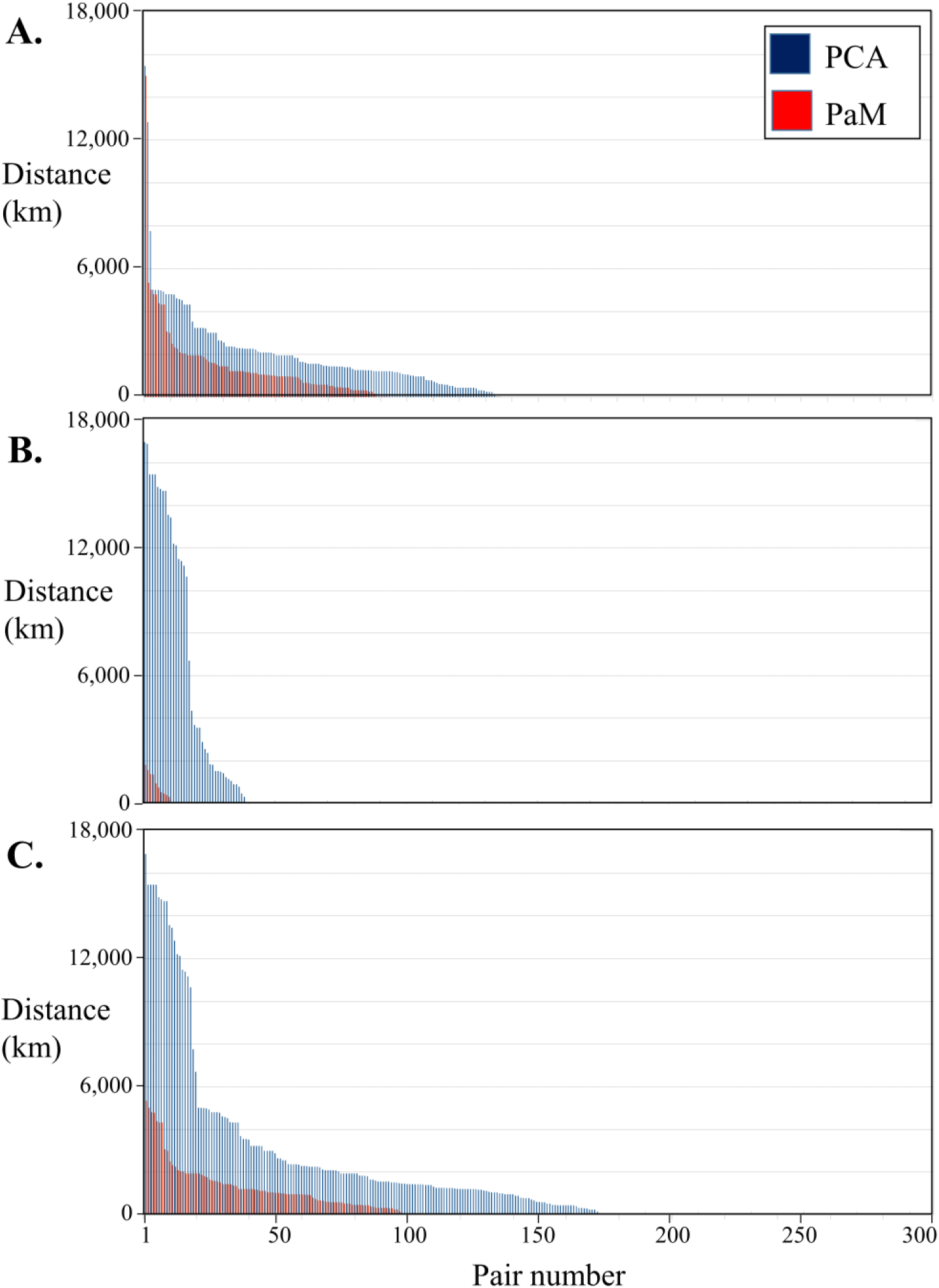
The geographical distance between individual pairs inferred by *PaM* and PCA. Geographic distances are calculated between pairs where both individuals are within the IBS-defined clusters (A), where individuals are in different clusters (B), and for all individuals regardless of cluster assignment (C).

We last compared the accuracy of *PaM* and PCA in identifying 13 mixed Indo-British individuals who formed a part of the Tartar/Tajikistan IBS cluster (Figure 6). PCA paired the Indo-British individuals with people from Tajikistan (4), Iran (2), Tatar (1), Russia (1), Ingush (1) and India (1). It correctly made one Indo-British pair and left out one individual. By contrast, *PaM* formed 6 Indo-British pairings, leaving the 13^th^ individual unpaired (although the Indo-British were part of the same IBS cluster consisting of Tartars and Tajikistanians). These results highlight the accuracy of *PaM* and its ability to identify and pair mixed individuals as compared to PCA and IBS.

### Running time

Running on a single core Intel i5 computer, *PaM*_*simple*_ (with threshold) finds the near optimum pairings in ~15 minutes for a test cohort of 1000 individuals, whereas *PaM*_*full*_ (with threshold) finds the optimised pairings in ~3 hours. If accessed online, results are typically emailed within 20 minutes.

### *Software* availability

*PaM* is freely available as a downloadable R package from https://github.com/eelhaik/PAM (16 Mb). In addition, a web-service has been created that allows users to upload genetic and demographic data for their test cohort and receive the optimised pairings solution by email (*www.elhaik-lab.group.shef.ac.uk/ElhaikLab/*).

## Discussion

Clinical trials are required to determine drug performance on multiple cohorts of sizes ranging from 40 to 10,000, where participants are split into treatment and control groups. The outcomes of these trials determine whether a drug should be tested in a larger cohort and, if successful, approved for use (Roy 2012). To evaluate the therapeutic effects of the tested drug, the treatment and control groups have to be as genetically homogeneous as possible (Yusuf and Wittes 2016) to minimize the variation in the response that is due to different genetic backgrounds (Figure 1). For that reason, pairing of cohort individuals is typically done at random after controlling for demographic criteria (e.g., age, sex, and self-reported race) and *a priori* to the trial. However, randomisation does not cure *population stratification*, particularly in initial small cohorts. Population stratification leads to biased results that, if positive, may not be replicated in a follow up larger trial and, if negative, may disqualify an effective drug. Correcting for population stratification *posteriori* to the trial is also problematic due to the difficulty in modelling ancestry and admixture and the reliance on “self-defined” race, a highly unreliable predictor (De Bono 1996; McAuley et al. 1996; Fustinoni and Biller 2000). Here, we present Pair Matcher (*PaM*) tool, designed to addresses the stratification bias *a priori*- and\or *posteriori* to the trial by identifying homogeneous treatment-control pairs using both genetic and demographic data. We evaluated *PaM*’s performances on simulated and real datasets against assignments made at random, based on racial classification, and PCA. We showed that *PaM*_*simple*_ outperformed the alternative classifiers.

Since *PaM*_*simple*_ is unable to utilise unpaired individuals, it may result in suboptimal pairings. This issue was resolved by introducing a threshold for the minimum acceptable genetic similarity between tested pairs, which significantly reduced spurious assignments. Using *PaM*_*full*_, which performs a nearly exhaustive search, reduced miassignments up to a relatively high perturbation level beyond which individuals were still assigned to genetically similar pair, though not necessarily their original counterparts. Due to the long computational time of *PaM*_*full*_, we recommend using *PaM*_*simple*_ with a nominal threshold of 7.

Overall we showed that *PaM* employs both demographic and genetic criteria to find optimised or near-optimised pairings solution for test cohorts consisting of unmixed and mixed individuals. *PaM* admixture model has several more advantages: first, the admixture components are calculated relative to the putative ancestral populations so their meaning remains the same between different analyses, second the admixture components of the individuals can be used to infer their geographical origins and ancestry with tools like the Geographic Population Structure (GPS) (Elhaik et al. 2014), and finally third it allows characterizing subgroups of responders, which can promote personalised medicine solutions tailored to population groups. *PaM* can be accessed online or be installed on the local computer. We hope that *PaM* will become a viable tool for clinical trials and association studies.

## Methods

### Simulated population datasets

We generated 24 datasets that comprised of 980-1000 individuals in ADMIXTURE’s Q file format (individuals x proportion of admixture components) (Alexander, Novembre, and Lange 2009). Here and throughout this work, we adopted the admixture model of Elhaik et al. (2014) of nine admixture components representing: *North East Asia*, the *Mediterranean, South Africa, South West Asia, Native America, Oceania, South East Asia, Northern Europe*, and *Sub-Saharan Africa*. Each dataset consisted of a file with nine admixture components generated randomly for individuals and their matching pairs and normalized so that each row would sum to 1. Dataset 1 consisted of 500 identical pairs. The GD between the Datasets 2-8’s pairs was increased in a controlled manner by modifying the admixture components of one individual from each pair of Dataset 1 through a perturbation of X% subtracting from the odd numbered admixture components and added to the even numbered admixture components, where X values increase from 0 to 20%. The perturbation percentage was applied alternately (negative to the first component, positive to second component and so on) to prevent normalization to undo the perturbation.

To evaluate the pairing solution in less ideal datasets, the remaining datasets were created by removing random individuals from the original datasets. Datasets 9-16 were created by removing one individual from each cohort (*remove-1*), and Datasets 17-24 were created by removing 20 individuals (*remove-20*), leaving datasets of 999 and 980 individuals, respectively. As before, assignment accuracy was related to the correct pairing of the remaining individuals against their known pairs with some individuals expected to be unpaired due to the removal of their exact match.

### Real population dataset

We used the Genographic dataset (Elhaik et al. 2014) that comprises of ~128,000 markers genotyped on 633 unmixed worldwide individuals of known geographic origins. We also created 13 two-way mixed individuals by hybridizing 13 Indians with 13 British to yield a final cohort of 646 individuals. The hybridization was done by merging an even amount of random SNPs from random Indian and British individuals and calculating the admixture components of these genomes (Elhaik et al. 2014). The admixture components of the Genographic and simulated individuals (Elhaik et al. 2014, Figure 1) were provided as input to *PaM*.

### Comparing *PaM*_*simple*_ and *PaM*_*full*_

To calculate the genetic distance (GD) and the score, *PaM* analyses the age (optional), gender (optional), and admixture components for each individual in the studied cohort. These three parameters are obtained from PLINK’s fam file (Purcell et al. 2007) (using column 4 for age and 6 for gender) and ADMIXTURE’s *Q* file (Alexander, Novembre, and Lange 2009), respectively.

The GD between the paired individual is defined as 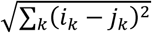, where *i, j* are the individuals with *k* admixture components. *PaM* calculates the *nxn* GD matrix for each possible pairing, where *n* is the number of individuals in the *Q* file. Each element of the matrix, specified by row *i* and column *j* corresponds to pair (*i, j*). The matrix is symmetric with respect to the diagonal, which contains all zeros. The corresponding *nxn* score matrix is calculated as follows: pairs that are age (within 5 years by default) and gender (mandatory) matched get one point. Nine potential points are awarded for every matching admixture component if |*ik*–*jk*| ≤1% for the pair (*i, j*). An ideally matched pair has a score of 10 (age\gender and nine admixture components). An ideal score for Datasets 1-8 is 5,000.

*PaM* operates in two modes: *PaM*_*simple*_ and *PaMfull.* The *PaM*_*simple*_ algorithm starts by selecting matrix row 1 (individual 1) and finding the column *j* which yields the minimum GD for pair (1, *j*). This matrix element corresponds to the first pair with row index 1 and column index *j*. Row 1 and column *j* and their symmetric element (*j*,1) are removed from the GD matrix. Row 2 is next selected (provided it has not been removed in the previous step) and the column which yields the minimum GD is selected to form the second pairing. The corresponding rows and columns are then removed from the matrix. The optimisation proceeds until all possible pairings are created and all unpaired individuals are stored. If the test cohort is an odd number then at least one unpaired individual is expected. The paired and unpaired individuals are reported in separate text files. To filter pairings with a score lower than a specified acceptable value, a threshold was implemented. A threshold of 7, for example, indicates that the pair’s age and gender matched as well as 6 of their admixture components.

In the case that matrix rows have multiple identical minima, there is a potential dilemma since the specific row minimum selection could affect subsequent pairings (due to the row/column removal upon pair selection), and the final pairing solution may not be optimal. We explored different selection schemes through exhaustive testing using single and random selection of the row minima, as well as a more complex method of minimising the sum of the remaining row minima, however the end results were very similar. Therefore, *PaM* uses a single minimum selection for each row, and this selection is the minima with the lowest column index *j*.

An extended version of *PaM*_*simple*_ is *PaM*_*full*_, which carries out a more exhaustive pairing search. *PaM*_*full*_ sorts in ascending order the test cohort data iteratively using the admixture components. The pairing procedure starts at a random row index (multiple times). The model starts by sorting the cohort data by the first admixture component then commences the search starting with a random row, *i*, index. The best pairing solution is stored. The cohort data is next sorted by the second admixture component, and the best pairing solution is found. If this solution minimizes the total GD of the final solution compared to the previous iteration then the ‘sorted admixture component 2’ solution is stored. The model proceeds by successively sorting the remaining admixture components to find the best pairing solution. Poor pairs are handled in a similar manner to *PaM*_*simple*_. However, when the data are re-ordered, all previously discarded individuals are included in the new solution search.

### Comparing *PaM* with demographic matches

*PaM* matches were compared with typical *a priori* matches based on age (within 5 years), gender, and “self-reported” race defined as “African,” “Asian,” “Latino” or “White.” Based on Elhaik et al.’s results (2014, Figure 1), ethnicity was inferred from the admixture components as follows: African ethnicity was assigned when the sum of *Sub-Saharan Africa* and *South Africa* admixture components was larger than 50%; *Asian* ethnicity was assigned when the *North East Asian* component was larger than 10%; and *Latino* ancestry was assigned when the *Native American* component was larger than 50%. All the remaining individuals were considered *White*.

Since “self-reported” race differs between studies, we considered three models: i) an individual is either African, Asian, Latino, or White. ii) an individual is considered either an African or non-African; iii) an individual is considered a mixture of Africans, Asians, Latinos, and Whites.

### Comparing *PaM* with PCA matches

*PaM* matches were compared with PCA-based ones (e.g., Luca et al. 2008). Briefly, PCA, converts a set of possibly correlated variables (here the alleles at each SNP) into a smaller set of linearly uncorrelated variables called principal components, or eigenvectors. Distances between individuals are determined by the major axes of variation in the eigenvector decomposition representation (EVD). The top *D* eigenvectors form a *D* dimensional map describing the ancestry of each individual, with the *d*th eigenvalue determining the importance of the *d*th dimension in the new representation of the data. Same-ancestry individuals have similar eigenvectors.

PCA was carried out using the SNP data using SNPRelate (Zheng et al. 2012). The EVD determines the distance between individuals on the basis of the top *D* eigenvectors serving as coordinates or dimensions and eigenvalues serving as weights to exaggerate differences in dimensions of greater importance. In all our analyses, the top two eigenvectors were used. These eigenvectors, calculated for each of the 646 individuals (Elhaik et al. 2014), were grouped into clusters of similar individuals. The clustering was achieved using the *k*-means clustering method *kmeans* in R. Similar to Luca et al. 2008, pairs were determined by a random assignment within each cluster.

To compare the quality of the results, the pairing solutions for *PaM* and PCA were evaluated by comparing the identity-by-state (IBS) clusters, inferred by PLINK (–cluster –matrix) (Purcell et al. 2007), and geographic distances, calculated using the Haversine formula (Gellert et al. 1989), for all the matched pairs. Due to the higher heterogeneity of the data and to reduce the number of unpaired individuals, a threshold of 5 was adopted for *PaM*.

## Authors’ contributions

E.E conceived the idea. D.R carried out all the analyses and developed *PaM*. E.E and D.R wrote the paper.

## Acknowledgment

E.E and D.R were partially supported by MRC Confidence in Concept Scheme award 2014-University of Sheffield to E.E. (Ref:MC_PC_14115). We thank Winston Hide for useful discussions.

## Supplementary materials

**Supplementary Table 1.** *PaM*_*simple*_ results without a threshold. Results are shown per dataset type and level of perturbation. The number of matched pairs are shown along with their total GD, total score, and the number of misassigned pairs. The remaining unpaired individuals are shown last.

|  | Perturbation (%) | Matched pairs |  |  |  | Number of unpaired individuals |
| --- | --- | --- | --- | --- | --- | --- |
|  |  | Number | Total GD | Total score | Misassigned |  |
| Full cohort | 0 | 500 | 0 | 5000 | 0 | 0 |
|  | 2 | 500 | 3.67 | 5000 | 0 | 0 |
|  | 5 | 500 | 9.13 | 4640 | 0 | 0 |
|  | 8 | 500 | 14.65 | 3319 | 0 | 0 |
|  | 11 | 500 | 20.21 | 2434 | 0 | 0 |
|  | 14 | 500 | 25.78 | 1952 | 6 | 0 |
|  | 17 | 500 | 31.91 | 1652 | 36 | 0 |
|  | 20 | 500 | 37.98 | 1496 | 73 | 0 |
| *Remove-1* | 0 | 499 | 0.102 | 4984 | 1 | 1 |
|  | 2 | 499 | 3.756 | 4984 | 1 | 1 |
|  | 5 | 499 | 9.442 | 4604 | 4 | 1 |
|  | 8 | 499 | 14.906 | 3300 | 4 | 1 |
|  | 11 | 499 | 20.326 | 2424 | 3 | 1 |
|  | 14 | 499 | 25.853 | 1941 | 9 | 1 |
|  | 17 | 499 | 32.096 | 1648 | 38 | 1 |
|  | 20 | 499 | 37.672 | 1496 | 64 | 1 |
|  | 0 | 490 | 3.778 | 4695 | 29 | 0 |
| *Remove-20* | 2 | 490 | 7.504 | 4663 | 33 | 0 |
|  | 5 | 490 | 12.888 | 4306 | 37 | 0 |
|  | 8 | 490 | 17.979 | 3119 | 39 | 0 |
|  | 11 | 490 | 22.652 | 2312 | 35 | 0 |
|  | 14 | 490 | 27.712 | 1874 | 38 | 0 |
|  | 17 | 490 | 33.622 | 1598 | 63 | 0 |
|  | 20 | 490 | 38.721 | 1452 | 99 | 0 |

**Supplementary Table 2.**
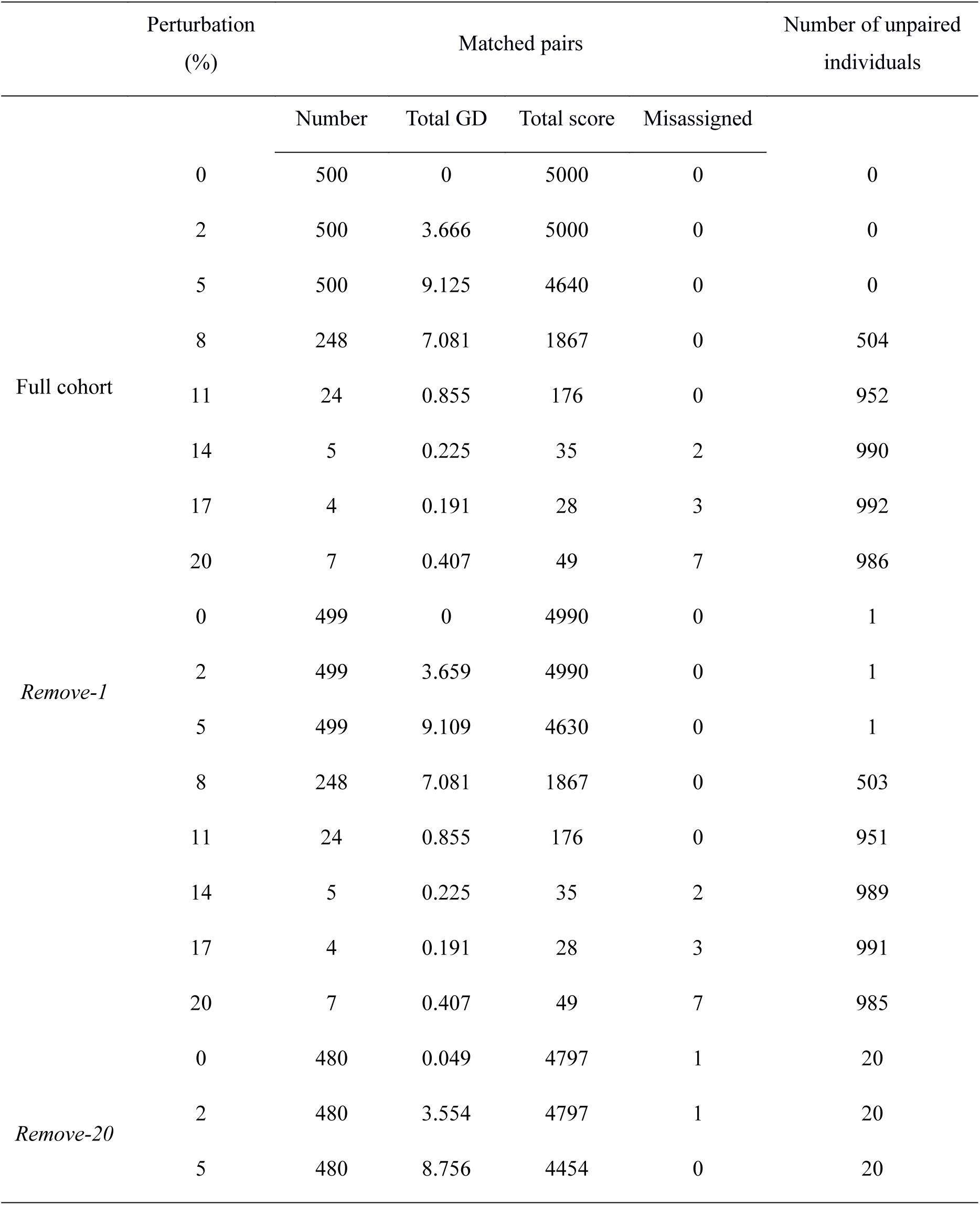
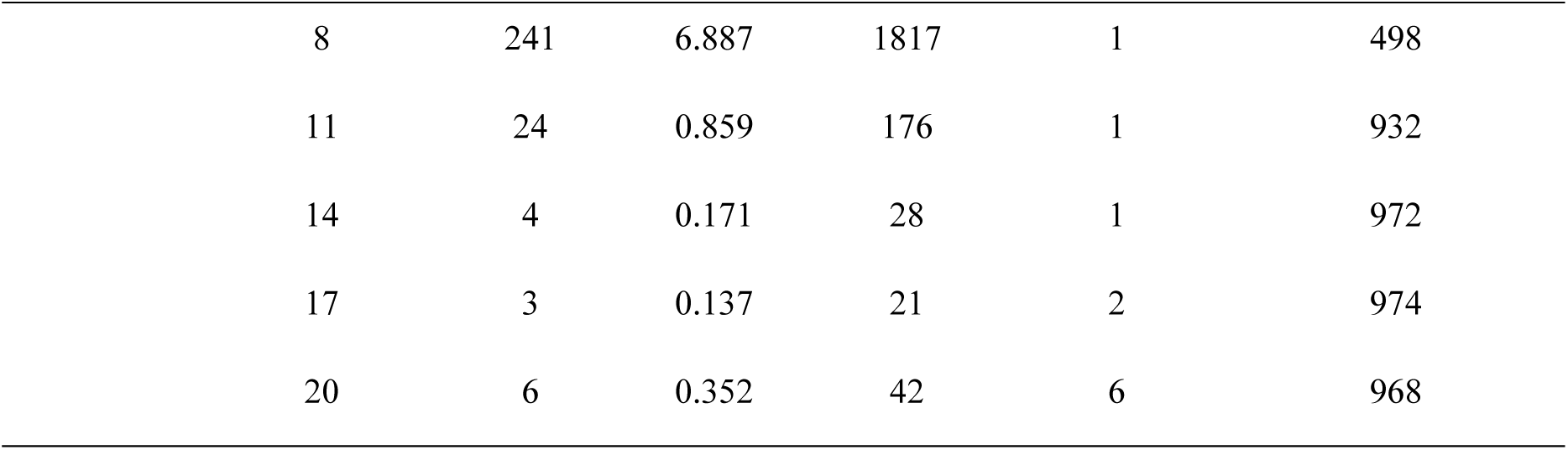
*PaM*_*simple*_ results with a threshold of 7. Results are shown per dataset type and level of perturbation. The number of matched pairs are shown along with their total GD, total score, and the number of misassigned pairs. The remaining unpaired individuals are shown last.

|  | Perturbation (%) | Matched pairs |  |  |  | Number of unpaired individuals |
| --- | --- | --- | --- | --- | --- | --- |
|  |  | Number | Total GD | Total score | Misassigned |  |
| Full cohort | 0 | 500 | 0 | 5000 | 0 | 0 |
|  | 2 | 500 | 3.666 | 5000 | 0 | 0 |
|  | 5 | 500 | 9.125 | 4640 | 0 | 0 |
|  | 8 | 248 | 7.081 | 1867 | 0 | 504 |
|  | 11 | 24 | 0.855 | 176 | 0 | 952 |
|  | 14 | 5 | 0.225 | 35 | 2 | 990 |
|  | 17 | 4 | 0.191 | 28 | 3 | 992 |
|  | 20 | 7 | 0.407 | 49 | 7 | 986 |
| *Remove-1* | 0 | 499 | 0 | 4990 | 0 | 1 |
|  | 2 | 499 | 3.659 | 4990 | 0 | 1 |
|  | 5 | 499 | 9.109 | 4630 | 0 | 1 |
|  | 8 | 248 | 7.081 | 1867 | 0 | 503 |
|  | 11 | 24 | 0.855 | 176 | 0 | 951 |
|  | 14 | 5 | 0.225 | 35 | 2 | 989 |
|  | 17 | 4 | 0.191 | 28 | 3 | 991 |
|  | 20 | 7 | 0.407 | 49 | 7 | 985 |
| *Remove-20* | 0 | 480 | 0.049 | 4797 | 1 | 20 |
|  | 2 | 480 | 3.554 | 4797 | 1 | 20 |
|  | 5 | 480 | 8.756 | 4454 | 0 | 20 |

|  |  |  |  |  |  |
| --- | --- | --- | --- | --- | --- |
| 8 | 241 | 6.887 | 1817 | 1 | 498 |
| 11 | 24 | 0.859 | 176 | 1 | 932 |
| 14 | 4 | 0.171 | 28 | 1 | 972 |
| 17 | 3 | 0.137 | 21 | 2 | 974 |
| 20 | 6 | 0.352 | 42 | 6 | 968 |

**Supplementary Table 3.** *PaM*_*full*_ results without a threshold. Results are shown per dataset type and level of perturbation. The number of matched pairs are shown along with their total GD, total score, and the number of misassigned pairs. The remaining unpaired individuals are shown last.

|  | Perturbation (%) | Matched pairs |  |  |  | Number of unpaired individuals |
| --- | --- | --- | --- | --- | --- | --- |
|  |  | Number | Total GD | Total score | Misassigned |  |
| Full cohort | 0 | 500 | 0 | 5000 | 0 | 0 |
|  | 2 | 500 | 3.666 | 5000 | 0 | 0 |
|  | 5 | 500 | 9.125 | 4640 | 0 | 0 |
|  | 8 | 500 | 14.645 | 3319 | 0 | 0 |
|  | 11 | 500 | 20.207 | 2434 | 0 | 0 |
|  | 14 | 500 | 25.781 | 1952 | 6 | 0 |
|  | 17 | 500 | 31.416 | 1633 | 21 | 0 |
|  | 20 | 500 | 37.023 | 1504 | 75 | 0 |
| *Remove-1* | 0 | 499 | 0.302 | 4978 | 2 | 1 |
|  | 2 | 499 | 3.991 | 4977 | 2 | 1 |
|  | 5 | 499 | 9.331 | 4619 | 2 | 1 |
|  | 8 | 499 | 14.809 | 3307 | 2 | 1 |
|  | 11 | 499 | 20.171 | 2429 | 0 | 1 |
|  | 14 | 499 | 25.74 | 1939 | 4 | 1 |
|  | 17 | 499 | 32.046 | 1630 | 33 | 1 |
|  | 20 | 499 | 37.427 | 1511 | 80 | 1 |
|  | 0 | 490 | 5.998 | 4555 | 50 | 0 |
|  | 2 | 490 | 8.482 | 4614 | 42 | 0 |
| *Remove-20* | 5 | 490 | 13.808 | 4261 | 45 | 0 |
|  | 8 | 490 | 17.948 | 3098 | 42 | 0 |
|  | 11 | 490 | 23.032 | 2309 | 38 | 0 |
|  | 14 | 490 | 27.852 | 1886 | 39 | 0 |
|  | 17 | 490 | 33.44 | 1598 | 67 | 0 |
|  | 20 | 490 | 38.441 | 1498 | 127 | 0 |

**Supplementary Table 4.** *PaM*_*full*_ results with a threshold of 7. Results are shown per dataset type and level of perturbation. The number of matched pairs are shown along with their total GD, total score, and the number of misassigned pairs. The remaining unpaired individuals are shown last.

|  | Perturbation (%) | Matched pairs |  |  |  | Number of unpaired individuals |
| --- | --- | --- | --- | --- | --- | --- |
|  |  | Number | Total GD | Total score | Misassigned |  |
| Full cohort | 0 | 500 | 0 | 5000 | 0 | 0 |
|  | 2 | 500 | 3.666 | 5000 | 0 | 0 |
|  | 5 | 500 | 9.125 | 4640 | 0 | 0 |
|  | 8 | 248 | 7.081 | 1867 | 0 | 504 |
|  | 11 | 24 | 0.855 | 176 | 0 | 952 |
|  | 14 | 5 | 0.225 | 35 | 2 | 990 |
|  | 17 | 4 | 0.191 | 28 | 3 | 992 |
|  | 20 | 7 | 0.407 | 49 | 7 | 986 |
| *Remove-1* | 0 | 499 | 0 | 4990 | 0 | 1 |
|  | 2 | 499 | 3.659 | 4990 | 0 | 1 |
|  | 5 | 499 | 9.109 | 4630 | 0 | 1 |
|  | 8 | 248 | 7.081 | 1867 | 0 | 503 |
|  | 11 | 24 | 0.855 | 176 | 0 | 951 |
|  | 14 | 5 | 0.225 | 35 | 2 | 989 |
|  | 17 | 4 | 0.191 | 28 | 3 | 991 |
|  | 20 | 7 | 0.407 | 49 | 7 | 985 |
|  | 0 | 480 | 0 | 4800 | 0 | 20 |
|  | 2 | 480 | 3.515 | 4800 | 0 | 20 |
| *Remove-20* | 5 | 480 | 8.756 | 4454 | 0 | 20 |
|  | 8 | 240 | 6.838 | 1810 | 0 | 500 |
|  | 11 | 24 | 0.859 | 176 | 1 | 932 |
|  | 14 | 4 | 0.171 | 28 | 1 | 972 |
|  | 17 | 3 | 0.137 | 21 | 2 | 974 |
|  | 20 | 6 | 0.352 | 42 | 6 | 968 |

